# Generalizable EEG encoding models with naturalistic audiovisual stimuli

**DOI:** 10.1101/2021.01.15.426856

**Authors:** Maansi Desai, Jade Holder, Cassandra Villarreal, Nat Clark, Liberty S. Hamilton

**Author notes:** **Corresponding Author:** Liberty S. Hamilton.

## Abstract

In natural conversations, listeners must attend to what others are saying while ignoring extraneous background sounds. Recent studies have used encoding models to predict electroencephalography (EEG) responses to speech in noise-free listening situations, sometimes referred to as “speech tracking” in EEG. Researchers have analyzed how speech tracking changes with different types of background noise. It is unclear, however, whether neural responses from noisy and naturalistic environments can be generalized to more controlled stimuli. If encoding models for noisy, naturalistic stimuli are generalizable to other tasks, this could aid in data collection from populations who may not tolerate listening to more controlled, less-engaging stimuli for long periods of time. We recorded non-invasive scalp EEG while participants listened to speech without noise and audiovisual speech stimuli containing overlapping speakers and background sounds. We fit multivariate temporal receptive field (mTRF) encoding models to predict EEG responses to pitch, the acoustic envelope, phonological features, and visual cues in both noise-free and noisy stimulus conditions. Our results suggested that neural responses to naturalistic stimuli were generalizable to more controlled data sets. EEG responses to speech in isolation were predicted accurately using phonological features alone, while responses to noisy speech were more accurate when including both phonological and acoustic features. These findings may inform basic science research on speech-in-noise processing. Ultimately, they may also provide insight into auditory processing in people who are hard of hearing, who use a combination of audio and visual cues to understand speech in the presence of noise.

**Significance Statement:** Understanding spoken language in natural environments requires listeners to parse acoustic and linguistic information in the presence of other distracting stimuli. However, most studies of auditory processing rely on highly controlled stimuli with no background noise, or with background noise inserted at specific times. Here, we compare models where EEG data are predicted based on a combination of acoustic, phonetic, and visual features in highly disparate stimuli – sentences from a speech corpus, and speech embedded within movie trailers. We show that modeling neural responses to highly noisy, audiovisual movies can uncover tuning for acoustic and phonetic information that generalizes to simpler stimuli typically used in sensory neuroscience experiments.

## Introduction

Humans live in a noisy world, where sound and speech perception often occur in the context of overlapping sounds. Understanding speech in natural environments involves the detection and parsing of acoustic and linguistic cues within these overlapping talkers and background noise. To understand how the brain performs this task, an increasing number of studies have attempted to address speech-in-noise perception by incorporating naturalistic stimuli in their experimental paradigms (Hamilton & Huth 2018, Fiedler et al. 2019). Previous findings have demonstrated that neural responses to naturalistic stimuli may actually yield comparable or better results to responses from a more controlled experimental paradigm (Hamilton & Huth, 2018, Huth et al. 2016, Wehbe et al. 2014, Lerner et al. 2011), and are more representative of our daily environment. What is considered “naturalistic” may vary: some studies use more naturalistic continuous or full sentence stimuli, while others use consonant-vowel (CV) syllables (Shankweiler & Studdert-Kennedy, 1966). Often the sentences used in these studies are presented in isolation and are far less “natural” than those used in natural communication. Many studies, including our own, have successfully used more controlled stimuli to model speech perception, such as sentences from the Texas Instruments Massachusetts Institute of Technology (TIMIT) corpus (Akbari et al. 2018, Hamilton et al. 2018, Tang et al. 2017, Chang et al. 2010, Mesgarani et al. 2014). Others have investigated EEG speech tracking using audiobooks (Broderick et al. 2019), which arguably are more natural than TIMIT sentences, but can lack the natural variation of pitch, timbre, other suprasegmental features of speech present in natural communication, and may often be read by only one talker. In addition, sentences from speech corpora like TIMIT are often repetitive and tedious to listen to for EEG tasks longer than one hour. Part of our motivation for this study was to utilize stimuli that were more engaging for participants and to investigate if neural responses can still be modeled robustly. A secondary aim was to investigate if neural responses and observed selectivity from the natural noisy contexts would generalize to responses to TIMIT, a noise-free and more controlled stimulus set.

Numerous EEG studies have demonstrated successful tracking of the acoustic envelope (Di Liberto et al. 2015, Fuglsang et al. 2017, Horton et al. 2011, Kubanek et al. 2013, Vanthornhout et al. 2018), phoneme and phonological features (Di Liberto et al. 2015, Di Liberto et al. 2019, Khalighinejad et al. 2017), pitch (Teoh et al. 2019, Krishnan et al. 2005), and even semantic information (Broderick et al. 2019) in speech. We expand upon these studies by investigating acoustic and linguistic feature encoding in both controlled and noisy, naturalistic stimuli. This allows us to assess the extent to which previous findings are stimulus-specific, versus which findings may generalize across very disparate stimulus sets.

Speech perception involves both auditory and visual domains, especially when a listener must comprehend speech in noisy environments. Multisensory integration has been particularly important in deciphering speech from noise, particularly for those with hearing impairments (Altieri & Wenger, 2013, Puschmann et al. 2019, Maglione et al. 2015, Hendrikse et al. 2019, Manfredi et al. 2018). Studies have investigated the influence of concurrent visual stimulation on auditory processing in the brain (Ozker, Yoshor, and Beauchamp 2018, Karas et al. 2019, Puschmann et al. 2019). These studies used stimuli in which audiovisual or various manipulations of audio or visual information could be used to determine what a given talker was saying. In Ozker et al., they used ECoG and fMRI to show that functional regions of pSTG were modulated by the addition of visual information congruent with auditory speech, and that this modulation was stronger when audiovisual speech was clear compared to when information in one modality was corrupted. Puschmann et al. were able to record EEG responses to naturalistic stimuli in individuals with high-frequency hearing loss due to presbycusis and found that increasing the use of visual stimuli compensated for the lack in auditory processing of speech in noisy environments. Both of these studies show that incorporating visual information, including lipreading, can enhance speech perception. While these results demonstrate that visual cues are important in allowing the listener to understand speech despite extraneous background sounds, it is unclear how other aspects of the visual scene that are not specific to a talker’s face may also modulate auditory signals.

In the current study, we used EEG to model neural responses to speech to two entirely different stimulus sets – controlled sentences from the TIMIT corpus, and audiovisual stimuli from children’s movie trailers. As mentioned previously, our goal was to determine to what extent previous findings of acoustic and phonological encoding in EEG are stimulus specific, versus which can be observed even in highly naturalistic, uncontrolled environments. One motivation was to quantitatively assess whether it is possible to replace some of the more monotonous stimulus sets for experiments where participants may not tolerate listening to these stimuli for a long time. In addition, by exploring how well encoding models trained on one stimulus set can generalize to another, in extremely different stimulus set, we can determine the robustness of our previous findings of acoustic and phonological encoding in the brain. To address these questions, our aims were to: 1) characterize neural responses to acoustic and linguistic features in the movie trailers versus TIMIT, 2) implement a cross-prediction analysis to identify if we could predict neural responses to clean speech from responses to speech-in-noise, and vice versa. 3) Since our movie trailer stimuli include visual components as well as auditory, we wanted to characterize how visual information influences auditory responses and the generalizability of linear models trained on multimodal stimuli. 4) Finally, because the movie trailers included time segments of clean speech and speech masked by noise, we wanted to identify if neural tracking of the noisy, naturalistic stimulus improved at specific times where speech occurs without any background sounds. Our results provide a framework for evaluating performance of models fit on naturalistic, noisy data, and show the extent to which these models generalize to a more controlled paradigm and vice versa. Finally, we demonstrate that visual and auditory information may be encoded separately for some stimuli, and that the influence of visual information on auditory input is likely stimulus specific.

## Materials and Methods

### Participants

Seventeen participants with typical hearing (8M, age 20-35, mean: 25.5±4.5 years) were recruited from the University of Texas at Austin. The ethnicity of our participants are as follows: 68% White, 13% Asian, 13% Hispanic, 6% African American. All participants were native English speakers. Pure tone and speech-in-noise hearing tests were performed using standard clinical procedures (ASHA, 2005) to ensure typical hearing ranges across all participants. Typical hearing responses for the pure tone test consisted of hearing thresholds <25 dB bilaterally for all frequency tones between 125 and 8000Hz, tested separately for each ear. Additionally, the QuickSIN test (Duncan and Aarts 2006) was administered to assess typical hearing in noise (no greater than 0-3 dB SNR loss). Participants provided their written consent for all portions of the experiment and were compensated at a rate of $15/hour for their participation. All experimental procedures were approved by the Institutional Review Board at the University of Texas at Austin.

### Stimuli and Task

Two contrasting stimulus types were used in this study. The first set consisted of sentence stimuli from the TIMIT corpus, which included continuous sentences spoken in English by multiple male and female talkers with no background noise or overlapping sounds (Garofolo et al. 1993). These stimuli also included transcriptions of the precise timing of the onset and offset of each phoneme and word. The second set of stimuli were children’s movie trailers, which contained overlapping speakers, music, and background noise (available at https://trailers.apple.com/). While these stimuli were entirely unrelated to the TIMIT sentences, members of the laboratory similarly transcribed the onset and offset of each word and phoneme, alongside a high-level description of the auditory environment (e.g. speech, speech with background noise, or background noise only) using ELAN transcription software followed by automatic alignment using FAVE-align, a modified version of the Penn Phonetics forced aligner (Rosenfelder et al. 2011). These timings were then manually corrected using Praat software (Boersma and Weenink). Each stimulus was transcribed by two authors (JH, CV, NC) to verify reliability of the transcribed boundaries. Although TIMIT and movie trailer stimuli were qualitatively very different in terms of the types of sounds present, we verified that the distribution of phoneme counts was comparable across TIMIT and movie trailers (2 sample Kolmogorov Smirnov test, D=0.1, *p*=0.99).

During the task, participants listened to 23 unique movie trailer stimuli alternated with TIMIT sentence stimuli in four blocks of 125 sentences each, and a fifth block of 100 sentences. The first four TIMIT blocks consisted of unique sentences with the exception of the final sentence. The fifth block contained 10 unique sentences with 10 repeats of each in a randomized order. For the movie trailer stimuli, 23 unique movie trailers were used, and each was presented once, however two unique stimuli (Inside Out and Paddington 2) were presented twice. The TIMIT sentences and movie trailers were presented through an iPad running custom software written in Swift (version 4, https://developer.apple.com/swift/), presented via an external monitor (see Data Acquisition).

During the task, participants were asked to watch and listen to the movie trailers but were not asked to attend to any particular speaker. For TIMIT, the participants were instructed to listen to the sentences while staring at a fixation cross. The overall task alternated between presenting five unique movie trailers and then 1 block of TIMIT sentences (125 sentences in the first four blocks and 100 in the fifth). This task format repeated a total of five times while giving the participants a short break in between. The duration of the task (excluding set up time) averaged at approximately one hour and a half, with about 40 minutes of TIMIT and 50 minutes of movie trailer data per participant. One participant (MT0007) was excluded due to poor quality of data.

### Data Acquisition

Neural responses were continuously recorded from a 64-electrode scalp EEG cap at a sampling rate of 25kHz using the BrainVision actiChamp system (Brain Products, Gilching, Germany). The impedance level for the EEG signal was kept below 15 kΩ. Additionally, eye movements were measured through electrooculography (EOG), with vEOG and hEOG measurements taken to aid in removing ocular artifacts from the neural data. The auditory stimuli were directly synchronized with the EEG data using a StimTrak stimulus processor (Brain Products). These stimuli were controlled by the experimenters outside of the EEG suite, with visual stimuli projected on a ViewPixx monitor inside the EEG suite. Audio levels were tested prior to the start of the task and were presented through insert earbuds (Etymotic, 2006) at a comfortable volume.

### Preprocessing

EEG and EOG data were downsampled to 128Hz using BrainVision Analyzer software. The remaining neural preprocessing steps were conducted using customized Python scripts and functions from the MNE-python software package (Gramfort et al. 2013). First, EEG data were re-referenced offline to the average of the mastoid electrodes (TP9 and TP10) and notch-filtered at 60Hz to remove any electrical artifact which may have been present in the EEG suite. Data were then bandpass filtered between 1 and 15 Hz using a zero-phase, non-causal bandpass FIR filter (Hamming window, 0.0194 passband ripple with 53 dB stopband attenuation, −6 dB falloff). This filtering approach has been used in previous studies of neural tracking of speech using EEG (Di Liberto et al. 2015, Broderick et al. 2019, O’Sullivan et al. 2014). Raw data were visually inspected and specific timepoints were manually rejected based on any non-biological sources of movement, such as if the participant moved or clenched their jaw and created electromyographic noise. No more than 10% of the data were manually rejected. An independent component analysis (ICA) was conducted to identify the components response for eye blinks and saccade artifacts as the EOG responses were recorded in separate channels alongside the 64-channels of scalp EEG data. Components reflecting ocular movements were subsequently removed from the data.

EEG data were epoched according to the onset and offset of acoustic stimuli to analyze the EEG signals which corresponded with the speech and speech-in-noise stimuli. The onset of each trial was identified through a customized script using a match filter procedure (Turin 1960), where the sound waveform of individual stimuli was convolved with the audio signal recorded to the EEG system, and the peak of the convolution was used to determine the offset and onset of each trial. Once data were epoched according to specific sentences or movie trailer stimuli, we then used the stimulus transcription files to identify the timing of specific auditory and visual features.

### Auditory and visual feature extraction

The auditory features extracted from our stimuli included phonological features, the acoustic envelope, and the pitch of each stimulus. For the phonological features, we created a binary phoneme feature matrix to indicate the timing of place and manner of articulation features for all phonemes for a given TIMIT sentence or for the movie trailer. Each element of the matrix was labeled with a 1 for the presence of a feature, and 0 for the absence of a given feature (Hamilton et al. 2018). Previous work using electrocorticography has demonstrated that neural responses across the STG respond to phonological features as opposed to the phonemes alone (Mesgarani et al. 2014). Researchers have demonstrated that these features are well tracked in EEG data as well (Khalighinejad et al. 2017, DiLiberto et al. 2015). Thus, we included the following place and manner of articulation features into the binary feature matrix: sonorant, voiced, obstruent, back, front, low, high, dorsal, coronal, labial, syllabic, plosive, fricative, and nasal. For example, the feature matrix would include a value of “1” in the “obstruent”, “fricative”, and “voiced” categories to indicate the onset of a /v/ sound.

The acoustic envelope of each speech stimulus was extracted using the Hilbert transform followed by a lowpass filter (3^rd^ order Butterworth filter, cut off frequency 25 Hz). The envelope, which represents the dynamic temporal changes in speech (Raphael et al. 2007), was extracted for each of the individual TIMIT and movie trailer audio files. Prior research has also shown that the auditory cortex tracks the pitch of a given sound (Chung and Bidelman, 2016, Tang et al. 2017, Teoh et al. 2019). Thus, we also included this feature in the model by computing the absolute pitch of each stimulus using the PraatIO package in Python (Jadoul et al. 2018), which provides a Python-based interface to the linguistics software, Praat.

A major difference between the TIMIT stimuli and movie trailer stimuli is the presence of visual information in the movie clips. Since concurrent visual information can also affect encoding of auditory features (Beauchamp, MS., 2005, Schneider et al. 2008, Holcomb et al. 2005, Kaiser et al. 2005, Molholm et al. 2004, Besle et al. 2009, Puschmann et al. 2019, Atilgan et al. 2018, Baskent & Bazo, 2011, Besle et al. 2008, Chandrasekaran et al. 2009, Crosse et al. 2015, Grant & Seitz, 2000, Kayser et al. 2010), we wanted to control for this potential difference in modeling EEG responses to our auditory features. This analysis also relates to our overall goal of understanding how generalizable encoding models can be when derived from much noisier, naturalistic stimulus sets as compared to TIMIT. Visual features were calculated for the movie trailer condition using a nonlinear Gabor motion energy filter bank (Nishimoto et al. 2011). Briefly, each frame of the movie was zero-padded with black pixels at the top and bottom (to convert a 720 x 1280 pixel frame into 1280 x 1280 pixels), and then each image frame was downsampled to 96 x 96 pixels. These frames were then converted to grayscale by first transforming RGB pixel values into L*A*B* color space and retaining only the luminance channel. Next, each grayscale movie was decomposed into 2,139 3D Gabor wavelet filters. These filters are created by multiplying a 3D spatiotemporal sinusoid by a 3D spatiotemporal Gaussian envelope. We used filters with 5 spatial frequencies, log-spaced from 1.5 to 24 cycles/image, 3 temporal frequencies (0, 1.33, and 2.667 Hz), and eight directions (0 – 315 degrees in 45° steps). Velocities were calculated over 10 frames of the movie at a time. We also included zero temporal frequency filters at 0, 45, 90, and 135 degrees, and one zero spatial frequency filter. The filters are positioned on each frame of the movie. Adjacent Gabor wavelets were separated by 4 standard deviations of the spatial Gaussian envelope. Each filter was also computed at two quadratic phases (0 and 90 degrees), as in Nishimoto et al. 2011. The Gabor features were then log-transformed to scale down very large values. Finally, we took the first 10 principal components of this stimulus matrix to reduce the dimensionality of the Gabor basis function matrix. Reducing the dimensionality from 2,139 to 10 PCs explained approximately 60% of the variance in the data.

This also corresponded to the point at which the second derivative of the variance explained curve approached zero (the “elbow” of the curve).

By incorporating these visual features into our model, we were able to regress out any EEG activity related to both static and moving aspects of the visual stimulus in the movie trailer stimuli. This also allowed us to assess whether including visual feature information significantly changes the measured auditory feature encoding weights.

### Encoding Models for Neural Tracking of Acoustic, Linguistic, and Visual Features

To model EEG responses to both audio and audiovisual stimuli, we used forward modeling TRF linear regression with different sets of acoustic, linguistic, and/or visual features. The multivariate temporal receptive field (mTRF) approach is used to describe the selectivity of neural responses to a given stimulus (DiLiberto et al. 2015, Crosse et al. 2016, Hamilton et al. 2018, Mesgarani et al. 2014, Theunissen et al. 2000). The goal of the forward modeling TRFs is to describe the statistical relationship between the input (auditory speech feature or visual feature) and output (the predicted EEG response based on the stimulus features). All 64 channels were used for all EEG participants, and separate models were fit to predict the activity in each EEG channel. The equation for the forward model TRF is shown below:

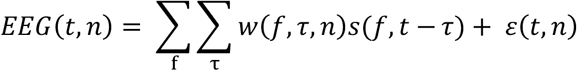

This model calculates the instantaneous neural response *EEG* at time *t* from electrode *n* and is expressed as a convolution between an input speech stimulus property, *s*(*f*, *t* – *τ*), with the EEG TRF weights, *w*(*f*, *τ*, *n*). The TRF is thus the mathematical transformation of a stimulus feature *f* into the EEG signal at different time lags *τ*. 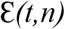 are the residual values from the linear regression otherwise not accounted for the input values. Figure 1 depicts all components of the forward modeling paradigm. The schematic for the predicted response from the TRF at a specific channel can be seen for the three unique features (pitch, acoustic envelope, and phonological features) as well as the combined model, which incorporates all three unique features as individual (*f*) values into the TRF model. For our audiovisual analysis, we could also simultaneously model responses to auditory and visual features of the stimulus. This framework allowed us to test different hypotheses about which features (acoustic, linguistic, or visual) were represented in the EEG data, and whether we could model how the brain tracks these features in a highly uncontrolled and naturalistic stimulus that contains background noise, music, or overlapping speech, with comparable fidelity to speech in isolation. A schematic including visual features is shown in Figure 4.

**Figure 1:**
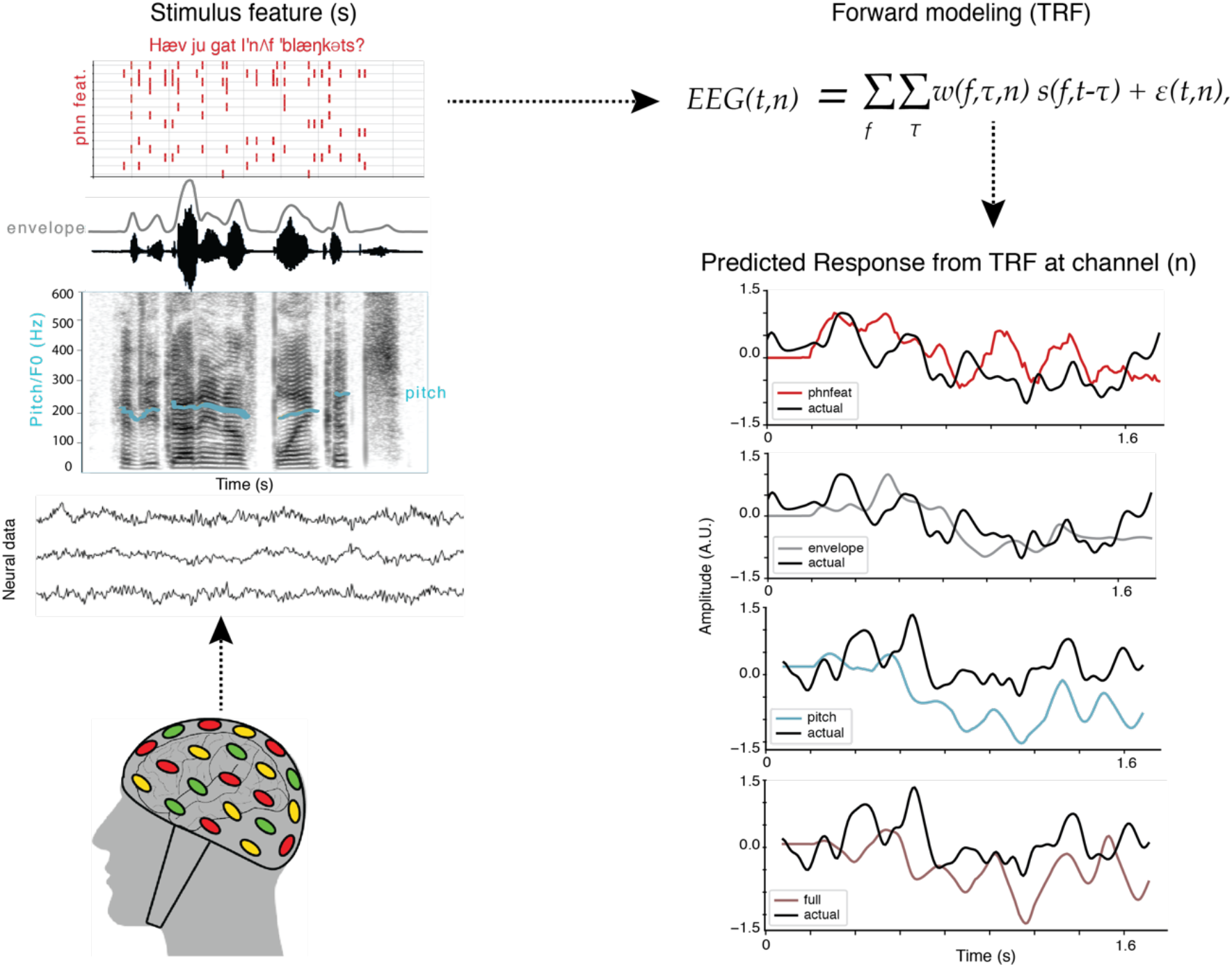
Analysis schematic showing how multivariate temporal receptive field (mTRF) models can be used to predict EEG responses to a given speech feature. We fit encoding models to neural data collected from participants as they listened to sentences from the TIMIT corpus and as they watched movie trailers, which contained speech in the presence of background noise. Identified speech features for analysis included the acoustic envelope, phonological features, pitch, and a combined or full model consisting of all the aforementioned features. Forward modeling was utilized to compute the mTRF, which was then used to predict the neural response to a specific speech feature (pitch, acoustic envelope, phonological feature) or a combination of features from a given EEG channel in both conditions from the EEG task (TIMIT and movie trailers). Adapted from (Crosse et al. 2016).

For all acoustic and linguistic feature models, we fit multivariate TRFs (mTRFs) using time delays from 0 to 600ms, which encompasses the temporal integration times for such responses as found in prior work (Hamilton et al. 2018). The weights (*w*) were fit using ridge regression on a subset of the data (the training set), with the regularization parameter chosen by a bootstrap procedure (n=100 bootstraps) and tested on a validation set that was separate from our final test set. The ridge parameter was chosen as the value that resulted in the highest average correlation performance across all bootstraps, and was set to the same value across electrodes. We tested ridge parameters of 0 as well as from 10^2^ to 10^8^, in 20 log-spaced steps. For TIMIT, the training set consisted of 489 out of 499 sentences (the TIMIT blocks 1-4, described above). The 10 unique sentences that were heard in TIMIT block 5 and were also heard in blocks 1-4 were used as the test set, so no identical sentences were used in training and testing. For movie trailers, the training set consisted of 21 out of the 23 movie trailers. The remaining two movie trailers were averaged and used as the test set evaluate the model performance was then used to evaluate the model performance in which these repetitions were averaged and used as the test set for the model for the clean speech condition. In a separate analysis, we also tested the effect on model performance when repeating the movie trailers in the test set up to 12 times.

In order to statistically evaluate the model performance in TIMIT, we conducted a Friedman ANOVA, which is a nonparametric version of the repeated measures ANOVA test. This was used rather than a standard ANOVA because our data violated the normality assumption. The independent variable was an individual feature whereas the dependent variable was the full model (a combination of all acoustic and linguistic features).

To compare performance across models, we plotted the individual model performance against the full model for each possible feature. To visualize the general distribution of correlation values, we plotted each point separately in addition to the convex hull surrounding those points (Figure 2). The significance of each model was determined by randomly shuffling the stimulus labels in 2-second chunks and computing a model based on this randomized data (using the ridge parameter from the true, unshuffled data), calculating the correlation between the predicted and held out data, and comparing that correlation to the unshuffled data. This was performed 100 times, which corresponded to a bootstrap p-value < 0.01.

**Figure 2:**
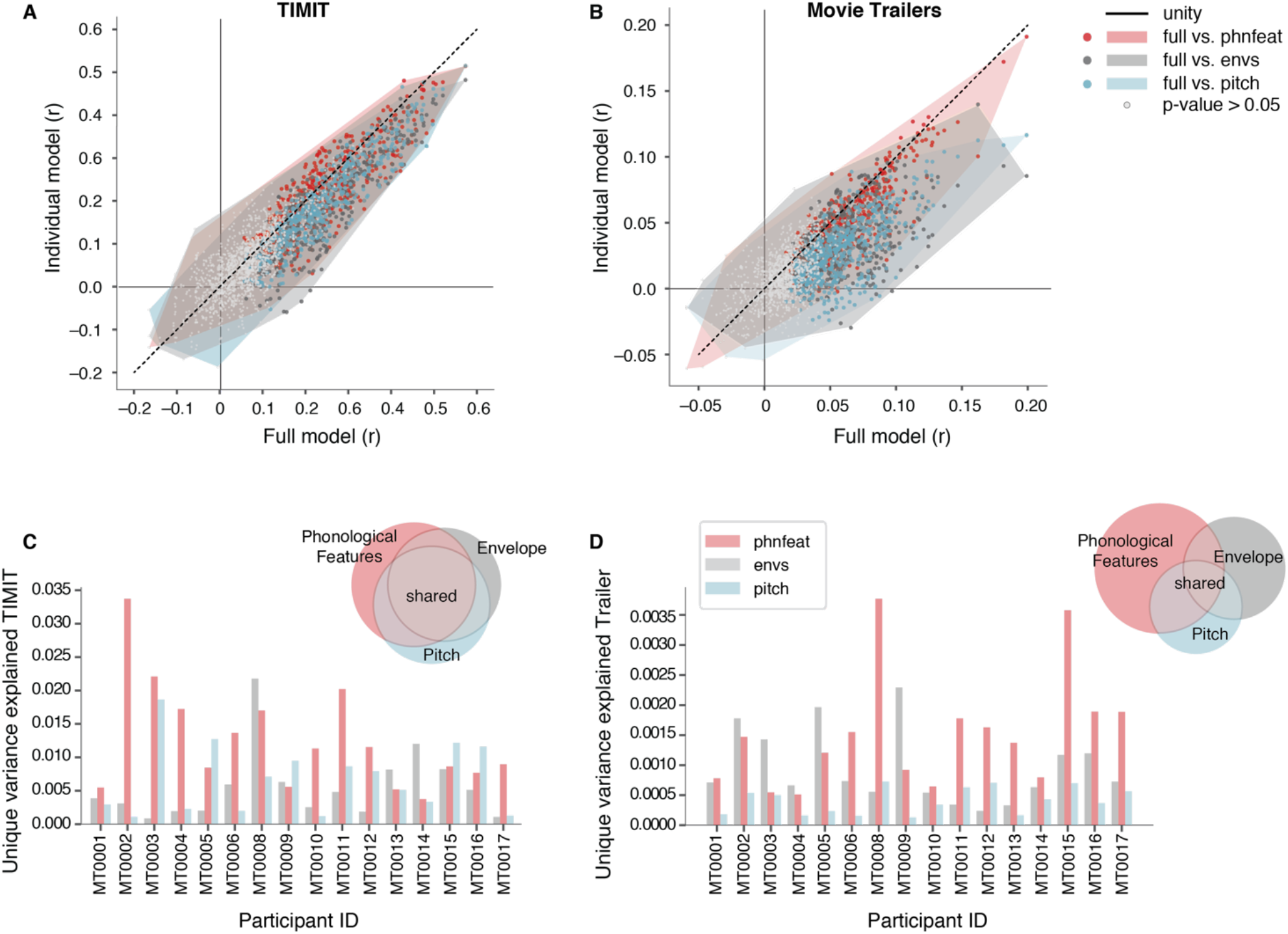
Contributions of phonological and acoustic representations in predicting EEG. Comparing individual speech features (pitch, acoustic envelope, phonological features) with the full model (combination of all auditory features). (A) Scatter plots showing the prediction performance (correlation between predicted and actual EEG data) of significant and non-significant electrodes for each condition in models fit using TIMIT responses. Each individual dot is a single electrode. Electrode color indicates the individual model type and the gray dots show non-significant electrodes across participants. The shaded regions indicate the convex hull around the scatter points for each comparison, to indicate how the points are distributed along, above, or below the unity line.(B) Same as (A), for movie trailers. (C) Variance partition analysis shows the average unique variance explained by individual features (phonological features, pitch, and envelope) for each participant separately (bar chart) and across all participants (pie chart) when fit on TIMIT data. (D) Same as C, for movie trailers condition.

### Variance Partitioning

A consequence of using natural speech (in both TIMIT and Movie Trailer stimuli) is that the stimulus features may be correlated with one another. For example, in the absence of background noise, the acoustic envelope is correlated with when phonological features occur in speech. To address this, we used variance partitioning to calculate the unique variance explained by each feature set. This represented the amount of additional variance that is added when including speech features in a model (de Heer et al. 2017). The purpose of this analysis was to identify the individual contribution of each feature, pairwise features, and full model to the overall model performance generated by the mTRF analysis. Furthermore, variance partitioning explains the intersection between multiple features. *R*^2^ values were calculated directly from the mTRF linear models. A total of seven unique features and intersections were used in this analysis.

The unique variance for a given feature was calculated by subtracting the R^2^ for a paired model from the R^2^ for a total model. For example, the unique variance for pitch was calculated by fitting the full model (envelope, phonological features, and pitch), and fitting a model with only envelope and phonological features. Taking the R^2^ value for the full model and subtracting the R^2^ value for the pairwise model would generate the unique contribution that was explained by adding pitch to the model. Unique individual features:

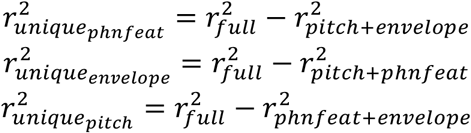

Equations for variation partitioning are shown below. In brief, we calculated model fits for each of the following features by using individual feature sets or the union of all pairwise and triplet combinations:

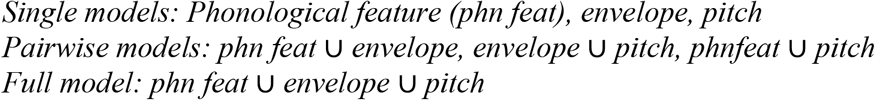

We then obtained the shared variance for each pair of models from the following equations:

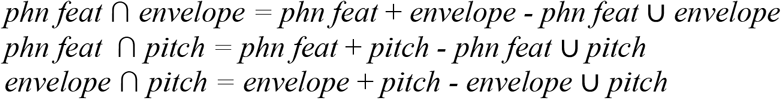

We then use these values to determine the intersection of all three models (the unique shared variance):

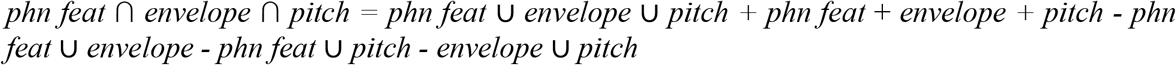

Finally, we could calculate the shared variance for each of the pairs of models without including the intersection of the combination of all three feature types:

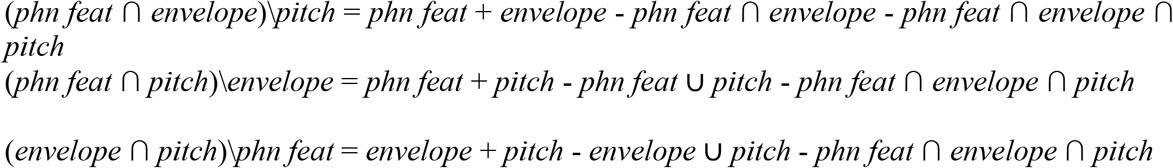

For some of the variance partition estimates, a correction needed to take place as certain models (single and pairwise) contained noise, some of which could be attributed to overfitting. In such cases, the variance partition calculations resulted in negative values where a post-hoc correction was used to remove the biased values (described in detail in DeHeer et al 2017).

### mTRF generalization across stimulus types

We next wished to assess the degree to which models trained on TIMIT sentences or movie trailers (MT) could generalize to the other stimulus set. This analysis allowed us to assess whether stimulus selectivity was similar across conditions, or whether neural tracking measures were stimulus specific. We conducted the same mTRF analysis for all speech features in both conditions from the EEG experiment to generate our weights and correlation values. We then used the weights calculated from using TIMIT as the training set to predict the neural responses of the respective speech features in the movie trailer EEG data. That is, using pretrained models from TIMIT, we then assessed model performance using movie trailers as the test set, rather than more TIMIT data. We then compared the correlation between the new prediction and actual EEG response to the correlation values generated by our original analysis, where training and test data came from the same stimulus type. We then performed the same analysis using pretrained models from the movie trailer data, evaluated on the TIMIT test set. The purpose of this analysis was to see if responses from TIMIT were generalizable to the responses from the movie trailers and vice versa. For example, to predict the EEG in response to the movie trailers from mTRFs calculated from TIMIT:

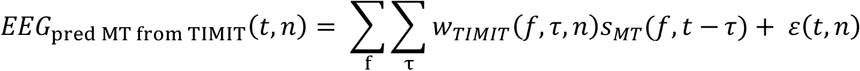

And to calculate EEG in response to TIMIT from mTRFs calculated from movie trailer stimuli:

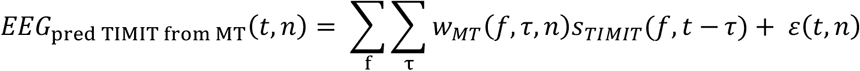

Finally, we determined whether there was a relationship between the performance on the cross-stimulus models and the original models by computing the linear correlation between the r values when the test set was held constant.

### Sliding correlation analysis

While the mTRF and the variance partition analysis demonstrated correlation values between the predicted and actual EEG and the unique feature contribution to model performance, respectively, they did not reveal whether specific time points within the stimulus were more reliably predicted by our model. We thus implemented a sliding window correlation analysis to assess the correlation between the predicted and actual EEG over smaller windows of time rather than over the entire test set. This procedure consisted of a defined sliding window time and identified the correlation coefficient between the predicted EEG data for a given feature, whether this was an individual feature (pitch, envelope, or phonological feature) or the combination of all three features. The equation is as follows:

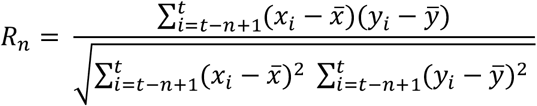

Here, *x* and *y* are the predicted and actual EEG, *t* is the time point within the sliding window, and *n* is the specified length of the sliding window to generate *R* at each time instance between the predicted and actual EEG for a given feature. We used a time window of size n=3 seconds, which is approximately the length of the longest TIMIT sentence.

To calculate how this sliding correlation varied based on the auditory environment, the movie trailers used in the test set were divided into three auditory environment conditions: speech alone, speech with background noise, and background only. The sliding window correlation was then averaged within each of the individual auditory environments for models using each speech feature (phonological feature, pitch, and envelope) in isolation and in the full model. We hypothesized that the phonological features during the clean speech only should generate higher correlation values overall for specific time points, some of which may be comparable to the correlation values seen in TIMIT.

### Effect of number of repeats of test set on model evaluation performance

The purpose of this analysis was to identify if increasing the number of repetitions for the movie trailer test set stimuli improved the overall model performance and thus, the correlation values between the predicted and actual EEG response. For our original analysis, the number of repetitions involved for the TIMIT and movie trailer test set stimuli varied based on stimulus condition. Out of the 499 TIMIT total sentences heard, 10 unique sentences were repeated a total of 10 times. The neural response to each unique sentence was then averaged and the aggregate of the averaged sentences were used as the test set. In contrast, out of the 23 movie trailers presented during the task, all of the trailers were heard once with the exception of *Paddington* and *Inside Out*, which were heard twice. The averaged neural responses for these two unique movie trailers were then used as the test set for the mTRF analysis.

To determine how this affected the observed model performance, we collected additional data from the same participant in two different sessions (MT0002 and MT0020). In the first session, MT0002 only heard two repetitions of the movie trailer test set stimuli alongside the remaining trailers. During the second session, the same participant (labeled as MT0020) heard the two unique test set stimuli a total of 10 times each, alongside the remaining trailers.

We performed the same mTRF analysis on the full model (containing all acoustic and linguistic features) in which we combined responses across both MT0002/MT0020 sessions to have a total of 12 repetitions for the test set stimuli in the movie trailers. The remaining 21 movie trailers responses were averaged and used as the training set. For each iteration, an additional repetition was added to a test set (eventually for a total of 12 repetitions) and the model was trained on the unseen movie trailers. This process was performed using a bootstrapping analysis such that different random subsets of 1 to 12 repetitions were used in the test set (100 bootstraps). This analysis examined how the model performance affected the average correlation value between the actual and predicted EEG data for each increase in test set stimuli in the movie trailer data set. Furthermore, we sought to identify if the average correlation values were comparable to those from TIMIT. To elaborate, do 12 repetitions of test set stimuli in the movie trailers have comparable model performance to 10 repetitions of test set sentences from TIMIT? Additionally, is there a specific point in the analysis in which adding more repetitions of the test set does not improve the average correlation from the model performance in each stimulus condition?

## Results

### Acoustic and linguistic representations of speech

Humans are rarely in an environment where speech occurs in the absence of any other background sounds. Still, many encoding model approaches rely on responses recorded in relatively controlled stimuli in the absence of noise, or in specific controlled noise conditions. While such models have been helpful in improving our understanding of the neural representations of natural speech, it is unclear whether these findings might generalize to a more naturalistic stimulus set, such as movie trailers. Thus, in our study we asked if we could identify which specific speech features were best tracked by neural responses in response to a more controlled continuous speech stimulus and a more naturalistic, multimodal stimulus. We also investigated whether comparable model performance was observed across both conditions, and whether this depended on the specific features included in the model. We chose the acoustic envelope, phonological features, pitch, and a combination of these three models to encompass a broad range of features that are involved in speech processing (Mesgarani et al. 2014, Oganian et al. 2019, Hamilton et al. 2018, Di Liberto et al. 2018). We trained on 64-channel EEG data responses using a full model (combining the acoustic and phonological features of envelope, phonological features, and pitch into one model) and individual speech features in both speech conditions (envelope only, phonological features only, or pitch only). Model performance was assessed by correlating the EEG responses predicted by our model and the actual EEG responses for held out data. We hypothesized that the combination of acoustic and linguistic features would contribute to higher model performance for movie trailers, since speech occurs in the presence of other unrelated sounds, and modeling these features separately can effectively lead to “denoising” of the data. In contrast, we expected that models based on predicting EEG from individual features (e.g. pitch or acoustic envelope or phonological features) should still perform relatively well when predicting responses to TIMIT. Thus, we first compared the correlation values between the individual models against the full model (Figure 2A-B). For both movie trailers and TIMIT, we found that the combination of acoustic and linguistic features yielded significantly higher model performance, as indicated by the majority of points falling below the unity line (*p*<0.001, Wilcoxon signed-rank test). Overall, the maximum observed statistically significant correlation value for the movie trailers (*r*=0.199, p<0.001 bootstrap test) was lower compared to TIMIT (*r*=0.57, p<0.001 bootstrap test).

In examining these conditions individually, we found that model performance across all participants in TIMIT for the individual features was more similar to the full model performance (Figure 2A). However, there was still a significant effect of model type when assessing correlation performance on TIMIT models (Friedman ANOVA Chi-sq=13.88, df=3, *p*=0.003). Post-hoc Wilcoxon signed rank tests were used to determine whether differences in each model type significantly differed, while controlling for within subject effects. These results showed that this effect was driven by significant differences between the full model and pitch (*p*=0.00089), along with the full model and the acoustic envelope (*p*=0.0098). In contrast, the post-hoc test showed that the performance of the full model and phonological features models were not statistically different from each other for TIMIT data (*p* = 0.22). Post-hoc tests demonstrated that, for TIMIT stimuli, correlations in the phonological feature model were not statistically different than the pitch model (*p*=0.24), the envelope model was statistically different than pitch (*p*=0.028), and that the statistical significance between the acoustic envelope and phonological feature models were at trend-level (*p*=0.057). Overall, these results demonstrate that the linguistic content from the phonological features seems to drive model performance and the addition of other acoustic features does not contribute to an increase in the correlation value.

To determine which features contributed significant unique variance in predicted responses to TIMIT, we used a variance partitioning analysis in which we were able to compute the unique variance explained by each individual feature, all pairwise combinations of features, and the variance shared by all acoustic and phonological features (Fig. 2C). The calculated variance partitions for TIMIT demonstrate more shared variance across individual features.

We next performed the same comparisons for models fit on the more natural movie trailer stimulus set, which included overlapping talkers and visual stimulation. We found a significant effect of model type across participants (Friedman ANOVA, Chi-sq=28.2, df=3, *p*=0.000003). Post-hoc Wilcoxon signed rank tests showed that the full model consistently outperformed the individual acoustic envelope (*p*=0.0027), pitch (*p*=0.0007), and phonological feature (*p*=0.0007) models. This indicates that including a combination of linguistic and acoustic information contributed to higher correlations in modeling neural responses to speech in this noisy listening situation (Fig. 2B). The variance partitioning analysis demonstrated that, unlike TIMIT, for movie trailer responses each individual feature contributed relatively more unique information to the overall model performance. That is, phonological features, pitch, and envelope were critical in modeling responses to the mixtures of sounds that occur in this more noisy, naturalistic stimulus. As with TIMIT, the phonological features contributed the most unique variance across participants (Fig. 2D). Unlike TIMIT, the amount of shared information was lower for the movie trailers compared to TIMIT, as indicated by a smaller relative area for shared variance in the Venn diagram. This suggests that, for a stimulus in which speech is accompanied by overlapping acoustics, neural tracking occurs in response to multiple individual features. While responses to linguistic information can still be modeled, we were also able to identify robust model performance for the acoustic envelope and additional unique variance explained by this feature. Much of this is likely attributed to the fact that the model is not only tracking the envelope of the speech, but also of the background sounds and music from the movie trailers. Overall, our results suggest that neural tracking for phonological features occurs both in noise-free as well as more uncontrolled, naturalistic settings despite the presence of varied background noise.

### Are receptive field models from each condition generalizable to the other?

Our previous analysis showed that we were able to predict EEG in noise-free continuous speech and in naturalistic, noisy conditions using linear models that incorporated acoustic and linguistic features. To extend on these findings, we asked whether models fit on one stimulus set would generalize so that they could predict neural responses for another stimulus type. If the neural responses could be generalized across stimulus sets, would it be possible to replace an experimental paradigm with stimuli that was more uncontrolled and naturalistic? To answer these questions, we used the weights from the individual feature models and the full model calculated from responses to TIMIT in order to predict responses to untrained TIMIT and movie trailer neural data. For each of these cross-predictions, we compare the performance of the model trained on one dataset and tested on the same stimulus type (e.g. predict TIMIT responses from TIMIT training data) or trained on a different stimulus type (e.g. predict TIMIT responses from movie trailer training data). This cross-prediction schematic is depicted in Figure 3A.

**Figure 3:**
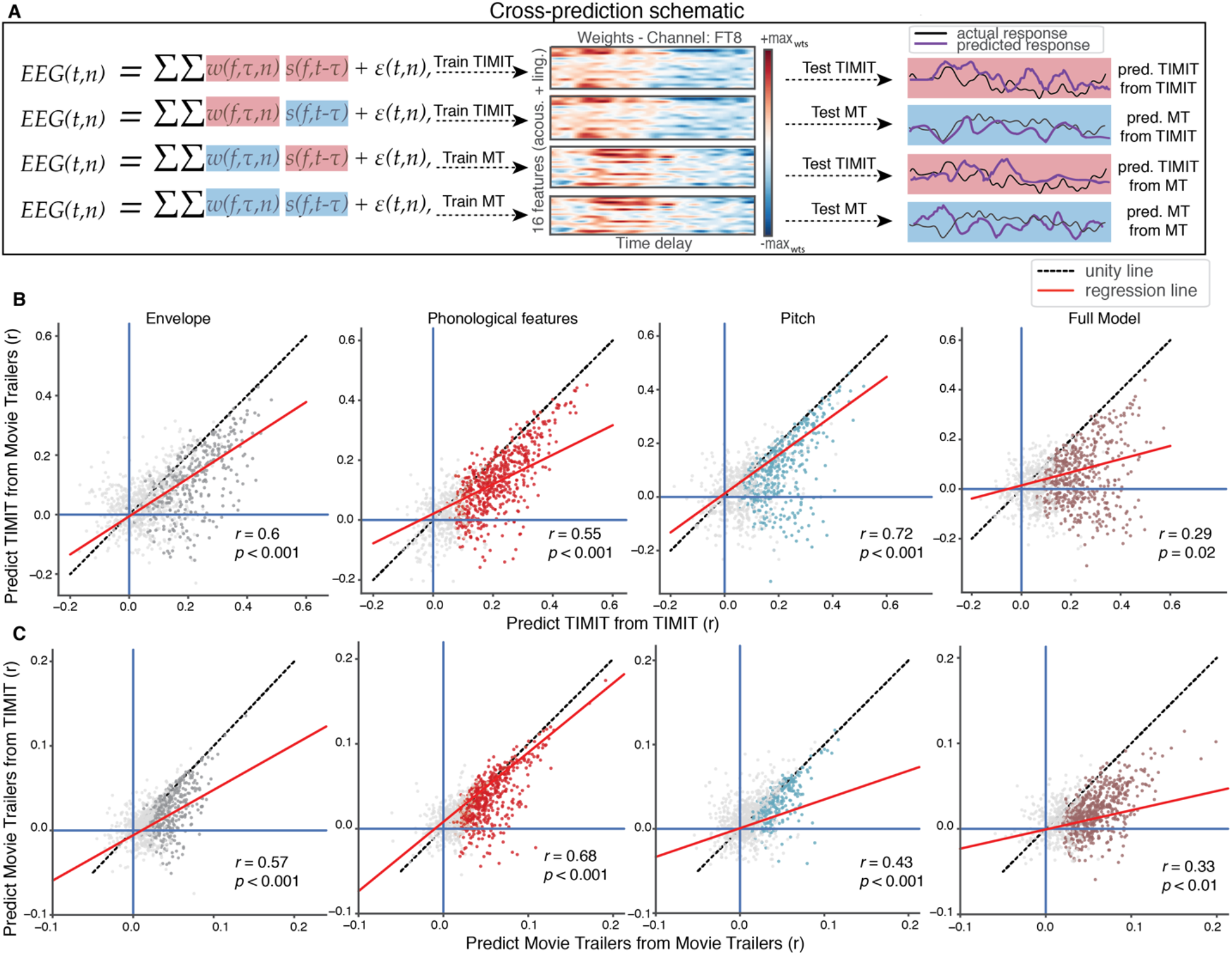
Cross prediction analysis shows that models fit to different stimuli are generalizable. (A) The cross-prediction analysis schematic is shown for the following conditions: training on TIMIT and testing on TIMIT, training on TIMIT and testing on MT, training on MT and testing on MT, training on MT and testing on TIMIT. In each of these training and testing conditions, the weights for the training condition were used for a given stimulus set and the stimulus feature for either the same or different stimulus was used for testing. The output generates a correlation value to assess model performance in how the new predicting for either the same or different stimulus condition. (B) Comparison of models trained on TIMIT data and tested on TIMIT data (x-axis) for all model types, both individual auditory features and a combined model. Using the mTRF weights derived from the opposite condition (movie trailers), we predicted responses to TIMIT (y-axis). Using the same features from either individual or the combined model, we predicted responses to movie trailers from the mTRF weights derived from TIMIT data. Despite relatively low prediction performance on the movie trailer data, the movie trailer trained models can predict responses to TIMIT. Each dot in the individual convex hull plots represents an individual electrode, where clusters of the same colored dots represent electrodes for a corresponding feature across participants. Dots in gray show correlations values where a bootstrap permutation test yielded p > 0.05, indicating that those electrodes with the respective r-values were not statistically significant for a given model. The unity line is shown as a dashed black line and the regression line alongside the corresponding statistic is shown in red in both (B) and (C). (C) Same as (B), but movie trailer individual or combined features are trained on movie trailers and tested on unseen movie trailer EEG data (x-axis).

We used the same convex-hull plots to show the correlation distributions for TIMIT and movie trailers. Correlation values below the unity line indicate electrodes for which the within-stimulus model performance was better, and above the unity line indicates better cross-stimulus performance. Overall, using the same stimulus type for training and testing (e.g. predict TIMIT from TIMIT or predict MT from MT) tends to result in better model performance. Specifically, there are more significant correlation values below the unity line. However, it was not the case that cross-stimulus predictions were invalid – in fact, the response to one stimulus could still be modeled from the other, although with slightly worse performance. We asked how correlated our model performance was when testing on a different stimulus type. If, for example, the same electrodes show good encoding model performance when tested on different stimulus sets, this is a good indication that our models are not predicting wildly different feature responses. Additionally, we wanted to examine the difference in generalizing stimuli when identifying individual feature types (e.g. pitch or envelope or phonological features) compared to the full model, which consists of the combination of acoustic and linguistic features. Across both conditions (Figure B-C), the results from the cross-prediction analysis demonstrated that the model performance comparing one condition (i.e. predict TIMIT from TIMIT or predict MT from MT) to the generalization condition (i.e. predict TIMIT from MT or predict MT from TIMIT) were statistically significantly related. This was the case across all feature condition types, whether this be the individual models or the full model. Interestingly, the cross-prediction analysis showed that the full model generally performed worse than the individual models (*r*=0.29, *p*=0.02 for predicting TIMIT from MT in Figure 3B and *r*=0.22, *p*=0.007 for predicting MT from TIMIT in Figure 3C).

Overall, we found that the model performance was better for TIMIT (Fig. 3B) when we trained and tested on TIMIT stimuli. While unsurprising, these responses were still able to be predicted from models using movie trailers as the training set, suggesting that feature encoding derived from neural responses to highly uncontrolled, noisy stimuli, can generalize to a more controlled stimulus set. The high model performance for predicting held out TIMIT EEG data from models also trained on TIMIT data could partially be attributed to the fact that listeners do not have to filter extraneous background sounds while attending to the primary speech source, so the signal-to-noise ratio of responses is higher. In addition, the correlation structure of the training/test stimuli is likely to play a role. Prediction correlations were generally lower for the movie trailers, but were still significantly higher than chance (Fig. 3C). While the model performance for the clean speech condition was better, we were still able to use the acoustic and linguistic feature models fit from the TIMIT training set and predict EEG responses to the movie trailers. Although these responses were shown to generalize to more a controlled stimulus set, such as TIMIT, the model performance correlations for movie trailers were still lower on average. Thus, we next examined whether these results were due to differences in the amount of testing data for both TIMIT and movie trailers, or whether this was due to inherent differences in the amount of speech alone versus speech in background noise.

### The effect of stimulus repetitions on model evaluation and performance

Our results in Figure 2 and Figure 3 demonstrated overall lower correlation values based on model performance for all feature types in the movie trailer condition compared to TIMIT. We were interested in whether and how the number of stimuli for the test set impacts the overall prediction performance from the model, and whether this differs for TIMIT versus movie trailers. We hypothesized that more repetitions of test set stimuli would improve the average correlation value, because averaging more EEG repetitions should increase the signal-to-noise (SNR) ratio of neural responses. We used the same TRF models in a single participant who had two separate recording sessions. In the first recording session the participant (identified as MT0002) listened to all blocks of TIMIT and listened to and watched all of the movie trailers. In this session, the participant watched and listened to two repetitions of the test set (*Paddington* and *Inside Out*) stimuli. In the second recording session, the same participant (MT0020) watched and listened to all of the movie trailers. However, during this session, MT0020 heard *Paddington* and *Inside Out* a total of 10 times each. No additional TIMIT sentences were played in the MT0020 session. We combined the test set stimuli from each recording sessions and found that there was an increase in average correlation as repetitions of the test set were added, with a maximum average value of *r*=0.079 (Fig 4A). A corresponding increase in model performance was also seen with additional repetitions of TIMIT sentences in the test set (Fig. 4B), however, the overall average correlations for this single participant were still greater than those of the movie trailers even with a small number of repeats. This suggests that the overall signal-to-noise ratio was not the only factor driving poorer model performance for movie trailers as compared to TIMIT. Including 10 repetitions of the movie trailer test set stimuli significantly improved the observed model performance values (Figure 4C-D, Wilcoxon rank-sum statistical test *Z* = 602.0, *p*=0.0034)

**Figure 4.**
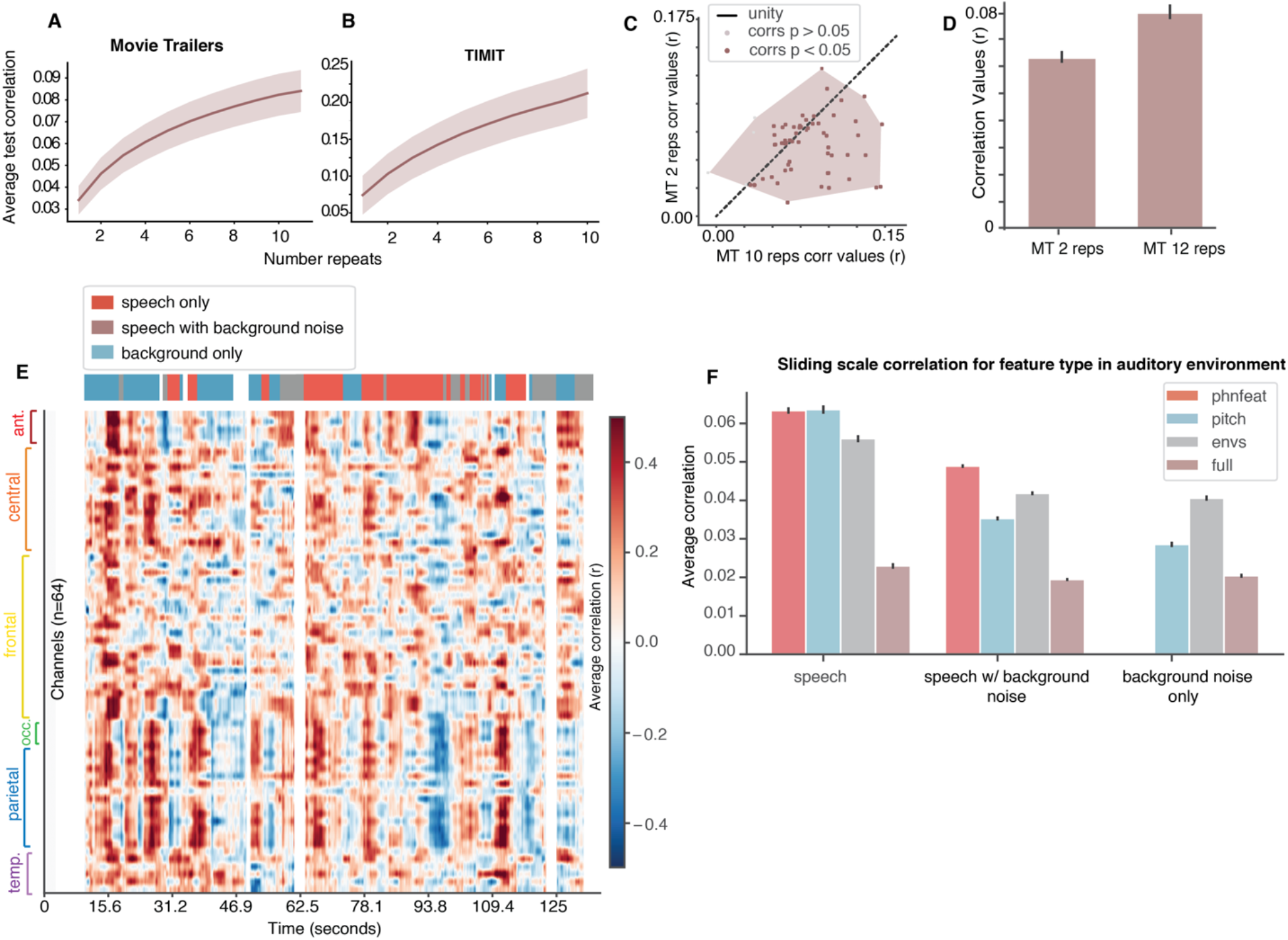
Evaluation of Movie Trailer prediction performance as a function of the number of repetitions in the validation set and as a function of time within the validation stimulus. (A) The average correlation value between the predicted and actual EEG data in response to movie trailers increases as repetitions are added to the test set. (B) Same as (A) for TIMIT. Note the overall higher correlation values compared to A. (C) Convex hull showing correlation values in one subject from two separate recording sessions, where the x-axis consists of 10 repetitions of the test set stimuli, whereas the correlations shown on the y-axis were evaluated using 2 repetitions of the test set. Each brown dot represents a single electrode. (D) Bar plot demonstrating the average correlation value improvement between each recording session (between 2 repetitions versus 10 repetitions of the test set stimuli), suggesting that the larger number of repetitions does generate a slightly higher correlation value on average (Wilcoxon sign-rank test, *p*=0.0034). Averages are across different partitions of the test set using a bootstrap procedure without replacement. (E) Top row shows auditory conditions within a single movie trailer (*Inside Out*) in which the overlapping auditory stimuli are separated based on speech only (red), speech with background noise (brown), and background noise alone (blue). This row is aligned with a heat map (below) from a sliding window correlation with a 3 second window for EEG responses predicted from phonological features from a single participant, demonstrating individual time points and channels that have higher correlation values based on the type of auditory stimuli occurring in the movie trailer. The y-axis contains the channels, divided based on the region of the EEG sensors (e.g. occipital, temporal, frontal). (F) Bar plot for the sliding window correlation across all participants (N=16) based on the auditory environment from both of the movie trailer test sets averaged together (Inside Out and Paddington) and separated based on an individual model (phonological features, pitch, acoustic envelope) or the full model (all individual features).

As the TIMIT condition consisted of continuous speech without any background noise, we hypothesized that prediction performance for the movie trailer dataset might be higher for time points where no background noise was present. Average correlation values were still considerably lower for more test set stimuli in the movie trailer condition, we hypothesized that the additional background noise in the test set stimuli could contribute to the lower correlation values. While some individual time points did have correlation values comparable to those of TIMIT (for example, the maximum correlations in Fig 4E are above r=0.4), this is not the case overall. Thus, we parsed the movie trailer testing stimuli into three different auditory environments: speech only, speech with background noise, and background only. We implemented a sliding window correlation analysis to investigate specific time points and channels in individual participants in each of the auditory environments. In Figure 4E, the heat map of the sliding window correlation for phonological features in a single participant shows that there are specific instances where it was possible to generate predictions up to *r*=0.4 over the 3-second time frame, as seen in times colored in dark red. Due to the 3-second sliding window time frame, it is possible to see that there is speech in the presence of background noise (e.g. at 62.5 seconds, the time point is blank demonstrating the lack of phonological features, but the legend above shows speech with background noise in brown). However, there were other time points where the model prediction generated negative values across channels, suggesting that attempting to predict EEG with phonological features was ineffective for some segments of the movie trailer. When examining the average correlation for these sliding windows within a particular stimulus segment across all participants (Fig. 4F), we observed the highest correlations for speech without background noise, followed by speech with background noise, and finally background noise alone. While the magnitude of these correlations was still lower than TIMIT, this suggests that the presence of background noise in the stimulus can also lead to less robust tracking of acoustic and phonetic features.

### Audiovisual components for speech tracking in a noisy environment

Up to this point, we have examined tracking of specific acoustic or phonological features in TIMIT and movie trailer stimuli. However, one obvious difference between the two stimuli is that the movie trailers also incorporated visual information, which has been shown to influence auditory perception (Beauchamp, MS., 2005, Schneider et al. 2008, Holcomb et al. 2005). If we wish to replace a more controlled stimulus like TIMIT with an uncontrolled stimulus like movie trailers, we must also examine to what extent the visual features influence auditory feature selectivity. As a final analysis, we sought to examine how the visual components were involved in speech tracking of the movie trailer stimuli. We built the same linear regression models, which now included all of the audio features (pitch, phonological features, and acoustic envelope) and added an additional set of visual features. The visual features were calculated from a motion energy model in which movie stimuli were decomposed into a set of spatiotemporal Gabor wavelet basis functions (Nishimoto et al. 2011).

These spatiotemporal Gabor wavelets allow us to investigate visual feature selectivity using Gabor wavelets that capture both static and moving visual aspects of the movie trailer. Fig. 5A illustrates some example spatiotemporal features, with each row representing a feature, and each column representing the evolution of that feature over time. The relative weights for each of these Gabor wavelets are used to construct a visual feature matrix (Fig. 5B) that describes visual motion parameters over time. In order to make the problem more tractable and to include a comparable number of auditory and visual features, we reduced the dimensionality of the Gabor feature matrix using a principal component analysis (PCA). A final example of the features that are used in the full audiovisual model is shown in Fig 5C, with some video stills to illustrate the visual scene at four example time points in our test set stimulus.

**Figure 5:**
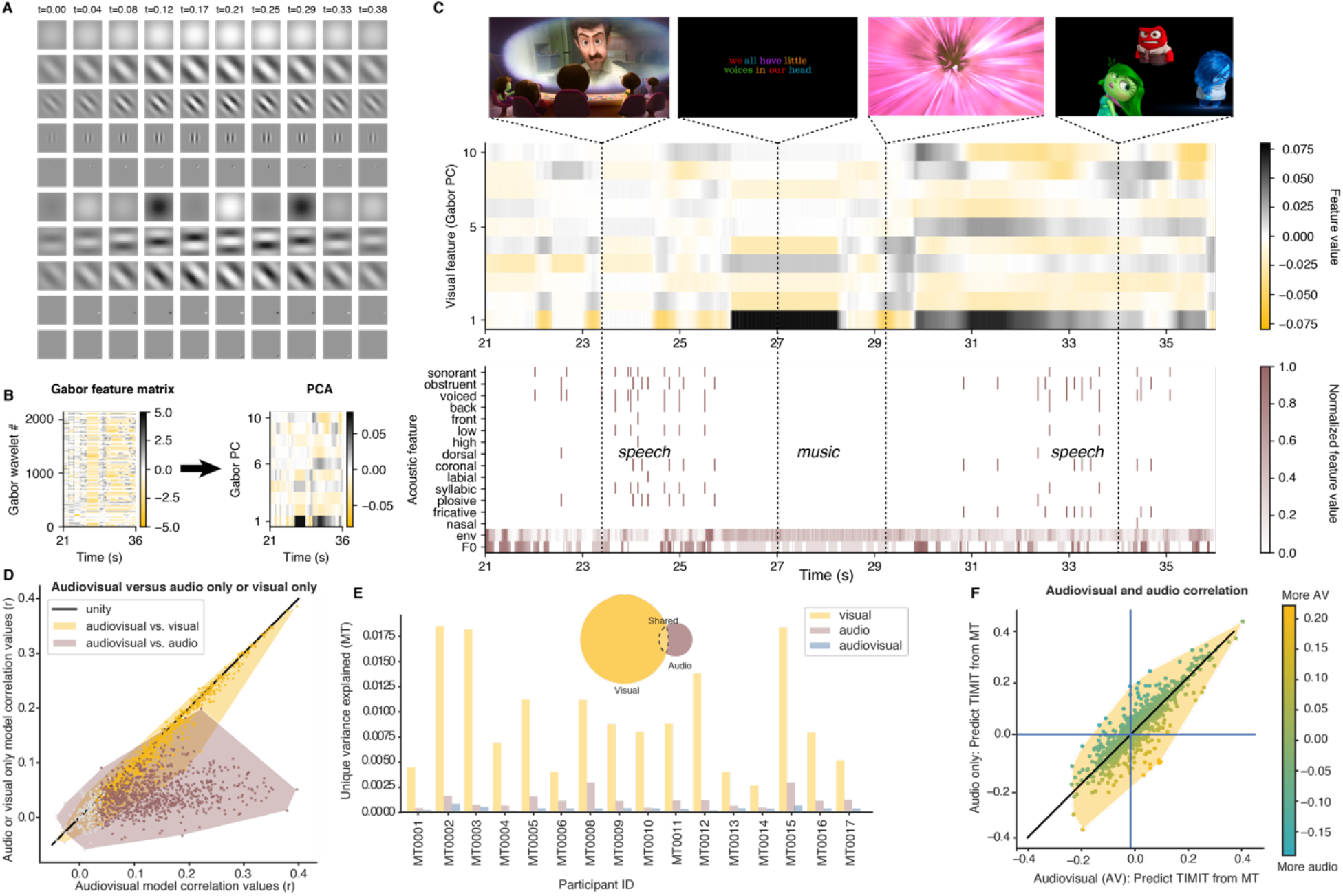
Unique contribution of visual features for the noisy (Movie Trailer condition) to assess model performance. (A) Visual stimuli in the movie trailers were decomposed into a set of Gabor wavelet features using a motion energy model. These features are essentially static or drifting gratings at different spatial and temporal frequencies. 10 example spatiotemporal Gabors are shown, where each row represents one spatiotemporal feature set, and each column represents the evolution of that feature over time. In our experiment, we used a total of 2,139 features, so these represent only a small fraction of the total set. (B) The 2,139 Gabor features are decomposed into their first 10 principal components using principal component analysis (PCA) on the entire Gabor feature matrix (2,139 features over time). For illustration purposes, only 15 seconds of data are shown. This reduced dimensionality matrix then serves as the visual input to our mTRF models. (C) Example combined visual and acoustic/linguistic features for 15 seconds of our test stimulus, “Inside Out”. The acoustic features are identical to those used in the previous model fits, while the visual features include the Gabors shown in (A-C). Example frames are shown for four timepoints in the stimulus. (D) Convex hull plots demonstrate model performance when comparing audio only responses to the combined audiovisual mTRFs along with the visual only responses. Each dot represents an individual EEG channel and the gray dots represent non-significant electrodes across both model comparison types. (E) bar plots demonstrate that the visual features heavily contribute to feature encoding in the noisy speech (movie trailer condition). Such results are further corroborated by the pie chart, demonstrating unique variance between the visual only, audio only, and shared audiovisual features. (F) A comparison of the cross-prediction analysis between the audiovisual stimuli in which the full model with visual features from the movie trailer condition were used to predict responses to the clean speech (TIMIT) condition, shown on the x-axis. The y-axis implements the same cross-prediction analysis but only uses the auditory features.

Next, we determined whether including visual features in the movie trailer model fitting influenced our ability to assess model performance between audio and visual only responses (Figure 5D). Adding additional auditory information alongside the visual, as depicted by the yellow region of the convex hull, improved overall model performance as compared to the visual-only responses (Wilcoxon rank-sum statistical test *Z* = 70.0, *p*<0.001). However, in contrast, when assessing the audio-only correlations from the model performance, adding visual information resulted in more robust model performance than just having the auditory features alone (Wilcoxon rank-sum statistical test *Z* = 110.0, *p*<0.001). We found that the visual stimulus information contributed a significant proportion of variance in the audiovisual speech condition (average unique max, *r*=0.4). In fact, the visual components occupied a significantly larger subspace in the bar plot compared to the audio only components (Fig. 5E). Despite this large contribution of visual information to overall explained variance, the shared region of the Venn diagram shows that each of these features (audio versus audiovisual) can be modeled separately, suggesting that the audiovisual stimulus does not significantly alter how the auditory features of speech are tracked in a noisy environment. To test this directly, we conducted a cross-prediction analysis (as in Fig. 3) to identify if we were able to predict neural responses to TIMIT from models fit using movie trailer training data with audio features only, and predict TIMIT from models fit using movie trailer data with the combined audiovisual features (Fig. 5F). Notably, no visual information was presented during TIMIT stimulus presentation, so the visual features are set to zero in the TIMIT stimulus matrix used for the prediction. We found that correlation values for both models were concentrated long the unity line, suggesting that the model performance was comparable for both conditions, and that regressing out the visual information did not significantly alter the predictability of the response to auditory information.

## Discussion

The use of naturalistic stimuli is becoming increasingly common in systems neuroscience (Hamilton and Huth 2018; Matusz et al. 2019). Understanding speech in real-world scenarios often involves parsing noisy mixtures of acoustic information that may include speech and non-speech sources. Natural environments are also inherently multisensory. Much of our understanding of how the brain processes speech relies on tightly controlled stimuli in the absence of noise, or parametrically controlled noise added to continuous speech stimuli. While these endeavors have been highly fruitful in uncovering tuning to specific acoustic and phonological features in speech, it is not clear to what extent models based on controlled stimuli can generalize to more complex stimuli. Taken further, in some experimental environments, it may be desirable to investigate speech processing using stimuli that are more enjoyable to listen to, so more data could be acquired without participants becoming bored or frustrated. Still, absent an evaluation of how such models generalize to more controlled datasets, it is difficult to interpret whether models based on naturalistic stimuli reflect the same processes involved in parsing more controlled stimuli. In this study, we addressed these questions by collecting neural responses to acoustic and linguistic features in clean speech (TIMIT) and naturalistic noisy speech (movie trailers) conditions using EEG. We were able to show that tracking of phonological features, pitch, and the acoustic envelope was both achievable and generalizable using a multisensory stimulus. In addition, despite the many differences between the TIMIT stimuli and movie trailers, we showed that models trained on the naturalistic stimulus could predict neural responses to controlled stimuli and vice versa, and that the visual stimulus, while a strong driver of neural activity, did not significantly affect encoding of auditory speech features.

In our first analysis, we described how individual acoustic or phonological features could predict brain data, and how that compared to a full model incorporating all of these features. While individuals have specific and unique abilities to track speech in noisy environments, we were still able to demonstrate that phonological features generally drove model performance for the clean speech condition. Generally, the acoustic envelope and phonological features were more predictive of EEG data than pitch, even though some of the statistical tests showed that the presence of pitch contributed to the full model in both conditions. A possible reason for lower correlation values in the individual pitch models in both speech conditions (TIMIT and movie trailers) could be attributed to the fact that pitch tracking may be more robustly identified in higher frequency EEG, such as tracking the FFR in the brainstem (Krishnan 1999, Galbraith et al. 2000, Zhu et al. 2013).

While our model performance correlation values were substantially lower in the noisy speech condition compared to the clean speech, it may be that the background noise corrupted measurement of the acoustic envelope from the individual sources, thus leading to lower performance when training the TRF model, predicting EEG responses to these features, and correlating these data with the actual EEG response from a given participant. Interestingly, this effect on the envelope may have been partially mitigated by including additional features for the pitch and phonological features in the movie trailer dataset, since the full model always outperformed any one feature set alone, and the shared variance was relatively low for this dataset (Fig. 2D). These stimuli involved a number of overlapping auditory stimuli (both speech and non-speech). As a result, additional information such as the acoustic envelope or even the pitch (while not as robust a model) could prevent one’s perception of the phonological features in the presence of such extraneous sounds. Furthermore, participants may have to pay more attention to the speech while deliberately ignoring all of the other sound sources. As demonstrated in this study, for the clean speech condition, the brain does not have to filter out extraneous sounds, which allows for better neural tracking of the given phonological speech target.

In addition to demonstrating neural tracking of acoustic and phonological features in the TIMIT and movie trailer stimuli, we were able to successfully perform a cross-prediction analysis, which suggests that the models are generalizable and not stimulus-specific. We demonstrated it was possible to use the weights from the movie trailer condition to predict responses to the TIMIT sentence, or a clean speech condition, suggesting that using brain data to model phonological and acoustic features in the presence of noise is possible, despite the highly noisy and uncontrolled stimulus. Additionally, the cross-prediction analysis alongside the variance partitioning suggested that the visual and audio components were modeled separately, and that the visual components did not affect the encoding processes in the brain for speech in noise in our dataset. While others have shown that visual information can significantly affect encoding of auditory information, it should be noted that much of the visual information in the movie trailers was not directly related to the speech. For a movie clip where much of the speech is accompanied by a view of the person talking (and their mouth moving), we might expect more overlap in the variance explained by auditory and visual features (O’Sullivan et al. 2017, Ozker et al. 2018, O’Sullivan et al. 2020). Future studies could investigate this directly by comparing movies where the auditory and visual information are either correlated (as in a “talking head” interview) or decorrelated (narration of a visual scene).

The sliding window correlation analysis demonstrated that there were specific times in the movie trailer data where the prediction performance from the linear regression model generated correlation values up to r=0.4 in an individual participant, comparable to those of TIMIT. However, when averaging the prediction performance across all participants across all timepoints for the speech only condition, the correlation values decreased substantially. The average of positive (upwards of *r*=0.4) and negative (minimum *r*=−0.4) correlation values may ultimately cancel each other out when analyzing the entire movie trailer. These lower values may be attributed to Simpson’s paradox, whereby model predictions within a sliding window may be unrelated to the prediction performance when a longer time window is included (Wagner 1982). In addition, when all participants are combined and the model prediction performance are averaged, differing SNR levels could also play a role in this result. As a future study, it may be useful to parse the speech and non-speech sentences and fit the same encoding models on multiple tiers of speakers and non-speech sounds in order to assess model performance.

## Conclusion

The current study contributes to the area of speech and speech-in-noise by investigating neural responses to acoustic and linguistic features and demonstrates that it is possible to predict neural responses from clean speech to noisy speech conditions, and vice versa. We found that higher-order linguistic cues drive neural responses when speech is presented in isolation. However, incorporating both acoustic and linguistic features are imperative in bolstering model performance in a highly naturalistic condition. Such results demonstrate that responses to these noisy and naturalistic datasets are generalizable to more controlled data sets, suggesting that we can utilize stimuli that may be both more representative of our daily environment and more enjoyable to listen to or watch. Our results also provide some intuition for how mTRF model performance changes based on stimulus characteristics as well as the amount of data and number of repetitions. Lastly, while audiovisual responses were greatly present our noisy, audiovisual speech condition, it was possible to separate neural responses to visual and auditory features. Such strategies could be incorporated into brain-based algorithms to assist cochlear implant or hearing aid users with improving speech in noise perception, as this is a typical complaint amongst individuals who present with hearing impairments and rely on amplification devices (Hagerman and Olofsson 2004, Nabelek et al. 1991, Levitt 2001).

## Author Contributions

MD and LSH conceived of the study and contributed to experimental design. JH, CV, and NC transcribed, aligned, and manually corrected annotations for the movie stimuli used in this study. MD, LSH, JH, CV, and NC collected the data. MD and LSH analyzed the data, prepared the figures, and wrote the manuscript. All authors read and approved of the manuscript.

## Acknowledgments

The authors would like to thank Garret Kurteff and Alexander Huth for helpful comments on the manuscript. The authors would like to Natalie Miller, Rachel Sorrells, Jacob Cheek, Garret Kurteff, and Ian Griffith for assistance in data collection. This work was supported by a grant from the Texas Speech-Language Hearing Foundation (The Elizabeth Wiig Doctoral Student Research Award, to MD).

